# High-resolution global recombination mapping in *C. elegans* reveals sexual dimorphisms shaped by meiotic chromosomal features and structures

**DOI:** 10.64898/2026.04.25.720830

**Authors:** Zachary D. Bush, John S. Conery, Hannah R. Wilson, Alice F. S. Naftaly, Devin Dinwiddie, Kenneth J. Hillers, Diana E. Libuda

**Affiliations:** Institute of Molecular Biology, Department of Biology, University of Oregon, 1229 Franklin Blvd Eugene, OR 97403, USA; Institute of Evolution and Ecology, Department of Biology, University of Oregon, 5289 University of Oregon Eugene, OR 97403, USA; Biological Sciences Department, California Polytechnic State University, San Luis Obispo, California, USA

## Abstract

Crossover recombination events during meiosis repair double-strand DNA breaks and ensure accurate chromosome segregation in most organisms. For many species, the genomic distribution of crossovers is nonrandom and sexually dimorphic. While many species evolved kilobase-scale “hotspots” for crossover formation, the *Caenorhabditis elegans* genome lacks hotspots, and crossovers are enriched across megabase-scale domains. Further, genetic and cytological studies indicate the crossover frequency in *C. elegans* spermatogenesis is higher relative to oogenesis in many but not all genetic intervals. To determine the genomic features that contribute to the sexually dimorphic recombination landscape in the absence of hot spots, we defined and analyzed the recombination landscape across the whole genome in *C. elegans* using whole-genome sequencing and high-resolution recombination mapping in single worms bearing recombinant chromosomes from individual sperm and oocytes. We find that the spatial distribution of crossovers is sexually dimorphic on chromosomes *I, II*, and *III*, and that the global rate of double-crossover events is 4.7-fold higher in spermatocytes. Additionally, we find that pairing and synapsis may contribute to the sexually dimorphic crossover landscape. In comparison to the spermatocyte crossover landscape, a higher proportion of oocyte crossovers are formed in the domains directly adjacent to the pairing centers of each chromosome. Further, reducing the genetic dosage of the synaptonemal complex central region protein SYP-2, which is a meiotic chromosome structural protein required for homologous chromosome synapsis, reshapes the oocyte crossover landscape to resemble observations in wild-type spermatocytes. Finally, we found that spermatocyte crossovers are partially enriched in H3K36me3-marked euchromatic regions, while many oocyte crossovers are enriched in H3K27me3-marked heterochromatic regions. Taken together, our studies reveal how synaptonemal complex component dosage and local chromatin states influence crossover placement and the sex-specific regulation of meiotic recombination.

**Author Summary:** Production of viable eggs and sperm depends on accurate chromosome segregation during meiosis. Segregation of parental copies of homologous chromosomes requires the reciprocal exchange and physical linkage of DNA that arises through crossover recombination. Increasing evidence indicates the existence of sexual dimorphisms during meiotic recombination. In this study, we generated and analyzed high-resolution recombination maps specific to spermatogenesis and oogenesis in the nematode *C. elegans*, which reveals sex-specific crossover distributions and a higher rate of crossing over in sperm cells. Further, we indicate how specific chromosomal features and structures differentially affect the crossover landscape in eggs versus sperm. Our work highlights how, in a system absent of pre-defined “hotspots” for recombination, local chromatin structures, chromosomal pairing domains, and the abundance of synaptonemal complex proteins are potential drivers for establishing the observable sex differences in crossover recombination.

## Introduction

Developing gametes, such as spermatocytes and oocytes, repair DNA double strand breaks (DSBs) with homologous recombination to form crossovers. Crossing over results in physical links, chiasmata, between the homologs and facilitates faithful chromosome segregation during meiotic cell division (1,2). A failure to induce or establish crossovers can lead to aneuploid gametes, infertility, and developmental disorders (3,4).

To ensure faithful inheritance of the genome, crossover formation is highly regulated in germ cells of many species. Since crossover formation is required for accurate homologous chromosome segregation in meioses of most organisms, the formation of at least one crossover between each pair of homologous chromosomes is ensured by crossover assurance and crossover homeostasis (5–10). DSBs, which are programmatically induced by the nuclease SPO11, are the initiating event for recombination (11,12), however, only a small subset of DSBs are repaired as crossover recombination events even in the event of an extreme excess of DSBs (8). Further, the genome-wide distribution of crossovers is non-random in most organisms. The formation of one crossover can inhibit the formation of crossovers nearby on the same chromosome through a phenomena known as crossover interference (13–19). Analyses of crossover distributions in many species reveals that the strength of crossover interference varies greatly between species and between sexes within species (20,21). In species such as *S. pombe* and *A. nidulans,* crossovers form with little to no interference (22,23), whereas some species such as *C. elegans* display nearly complete interference across the length of a chromosome and each set of paired homologs typically form only one crossover event (8,24). Previous studies in *C. elegans* have indicated that while chromosome structures such as the synaptonemal complex (SC), chromosome axis, and general chromatin condensation may regulate crossover placement and/or interference (25–28), there are likely additional multifaceted mechanisms regulating crossover rate and placement that remain to be uncovered.

Several studies indicate that biological sex can impact crossover regulation across the genome of many eukaryotes. Specifically, the rate and distribution of crossovers across the genome, known as the recombination landscape, often differ between oogenesis to spermatogenesis in multiple model systems. Evidence from studies in plants, mollusks, arthropods, amphibians, reptiles, birds, and mammals all demonstrate that the distribution of crossover events during spermatogenesis is slightly elevated in sub-telomeric regions in contrast to the more centrally located crossovers in oogenesis (29). Further, the crossover number per chromosome pair is also sexually dimorphic, with oogenesis exhibiting a higher crossover rate than spermatogenesis in many species (29).

Increasing evidence suggests chromosome structures can influence sex differences in the recombination landscape. Mammalian oocytes, but not spermatocytes, undergo global DNA demethylation (30,31), and DNA methylation has been shown to promote the initiation of recombination (32). DNA methylation occurs primarily at CpG nucleotides, which are enriched in sub-telomeric regions coincident with known male biases for crossing over (29,33–36). Additionally, studies in mice and plants have shown that chromatin modifications like H3K4me3 can differentially influence where recombination is initiated in each sex (32,37). Further, there is emerging evidence that sexual dimorphisms in meiotic chromosome structures, such as the synaptonemal complex (38), can contribute to sex-specific crossover patterning. Thus, sexual dimorphisms in meiotic chromosome structures and epigenetic modifications may be two of several factors underlying sexually dimorphic recombination landscapes.

Euchromatic regions of the genome, typically associated with transcriptional activity and accessible chromatin (39), may promote initiation, processing, and maturation of crossover recombination. Local chromatin structure and nucleosome positioning around promoter regions can influence the formation of programmed DSBs, the initiating event for homologous recombination, by the highly conserved, topoisomerase-like protein SPO11 (11,12,40–43). In mice and humans, the histone methyltransferase PRDM9 binds specific DNA motifs and creates “hotspots” for crossover formation (44–47). In species that lack PRDM9-mediated hotspots, such as budding yeast, DSBs and crossovers are enriched in physically accessible euchromatic regions, such as nucleosome-depleted sequences and gene promoters (48). In species that lack both hotspots and PRDM9, such as the nematode *Caenorhabditis elegans* and the fruit fly *Drosophila melanogaster*, local chromatin states are hypothesized to influence crossover placement. In fruit flies, both predictive models and experimental evidence points to a lack of recombination in heterochromatic regions of the oocyte genome (49–51). In *C. elegans* chromatin modifications such H3K9 methylation are known to play a role in shaping the recombination landscape in oocytes (52), and crossovers are preferentially positioned in multi-megabase domains at the terminal thirds of each chromosome (53,54).

*C. elegans* is an excellent model to identify hotspot-independent mechanisms that contribute to sexual dimorphisms in crossing over. Assessing crossovers at the resolution of multiple megabases via genetic assays has shown sex-specific differences in crossover frequencies on the arms of some autosomes (17,55–57). In *C. elegans* oocytes, crossover homeostasis and interference is incredibly robust, where paired homologs regularly receive only one crossover (8,24,25,58,59). In contrast, the incidence of double crossover events is slightly elevated in spermatogenesis across multiple genetic intervals (17,55,60–63), which is also supported in cytological studies by increased COSA-1 foci in sperm nuclei (64). This finding suggests that the strength of interference is likely sexually dimorphic and weaker in spermatogenesis. Notably, a few studies did detect double crossover events on some genetic intervals in oogenesis, but not spermatogenesis (17,56,62), thereby generating a debate on whether spermatogenesis indeed has a lower level of crossover interference in comparison to oogenesis. Further, the apparent absence of recombination hotspots in the *C. elegans* genome allows for the careful and unbiased dissection of which sequence-level, epigenetic, and/or higher-order chromosome structural features shape the recombination landscape and sexual dimorphisms therein.

Whole genome sequencing can be leveraged to illuminate both the crossover landscape and the genomic features that regulate sex-specific crossover positioning and distribution. Prior studies have mapped and examined finer-scale features of crossover recombination in oocytes and shown an association of the crossover rate near sites where homologous chromosomes pair (54). Additionally, fine-scale crossover mapping on a subset of the *X* chromosome in oocytes also showed a negative association of crossover sites with euchromatic histone modifications (65). Although the genomic landscape of chromatin modifications differs between *C. elegans* oogenesis and spermatogenesis (66), it is largely unclear whether these chromosomal features promote sex-specific crossover distributions in *C. elegans*.

To illuminate the both the fine-scale and broader scale genomic features contributing to the sexually dimorphic recombination landscapes in *C. elegans,* we performed high-resolution crossover mapping in individual products of both sperm and oocyte meioses. Using hundreds of thousands of SNP markers between the N2 Bristol and CB4856 Hawaiian strains, we mapped crossovers in spermatocytes and oocytes with an average resolution of one SNP every 300 bp. Our analysis demonstrates that Chromosomes *I, II,* and *III* display the most sex-specific differences in recombination rates at both the kilobase and megabase scales. The global rate of double crossover events is nearly five-fold higher in spermatogenesis than oogenesis. We find that perturbing the dosage of the synaptonemal complex central region protein, SYP-2, alters the crossover distribution and increases the genome-wide crossover rate in both oocytes and sperm. Finally, we find that the sex differences in the crossover landscapes are highly associated with specific chromatin states that regulate gene expression in the germline. Crossover formation in spermatogenesis is highly associated with the euchromatic histone modification H3K36me3, while oocytes crossovers display high association with the heterochromatic histone modification H3K27me3. Taken together, these results reveal that the mechanism(s) of homologous recombination that produce a sexually dimorphic crossover landscape in oogenesis and spermatogenesis are shaped by synaptonemal complex dosage and local chromatin states.

## Results

### High-resolution mapping of crossovers in single *C. elegans* genomes

To detect crossovers with high resolution in each sex, we performed whole-genome sequencing of individual F2 progeny harboring recombinant chromosomes from single meioses in the F1 generation. Briefly, F1 hybrid progeny were generated by crossing N2 Bristol hermaphrodites and CB4856 Hawaiian males. F1 progenies were then backcrossed to individuals with a Bristol genetic background so that F2 progeny inherit singular recombinant chromosomes from sperm or oocyte meioses along with another N2 Bristol homolog (Figure 1A). From the F2 generation, 300 oocyte-derived samples and 310 spermatocyte-derived samples were individually sequenced on Illumina’s NovaSeq platform with an average read depth of 10X across all SNP sites (see Methods; Supplemental Figure S1A). Using 213,591 SNPs identified between the N2 Bristol and the CB4856 Hawaiian strains (67), we were able to map crossovers (Figure 1B, for example) using our adaptation of the TIGER pipeline (Supplemental Figure S2) for HMM-based inference of crossover breakpoints, which enables robust crossover detection against samples even sequenced at low coverage (68). In our dataset of 610 genomes, our crossover mapping pipeline was able to successfully call crossovers in 297/300 oocyte samples and 300/310 spermatocyte samples. In the oocyte data, we detected 837 crossovers across 830 chromosomes, and 738 crossovers across 710 autosomes in the spermatocyte data (Table 1). The average size of the SNP intervals containing crossover breakpoints is 1,569 bp with a median length under 1 kb for each chromosome (Table 1), indicating great resolution of detection in both sexes. The increased number of detected crossovers relative to the number of unique chromosomes represented is due to the presence of double crossover events. On chromosome *I*, we detected approximately 23% fewer crossovers in oocyte samples. While some samples have insufficient read coverage and there are known genetic incompatibilities between the Bristol and Hawaiian backgrounds (69–71), chromosome *I* also displays a consistently distinctive pattern across multiple analyses in our dataset. Overall, our methods reliably detect crossover events on randomly segregating chromosomes through whole-genome sequencing of individual nematodes.

**Figure 1.**
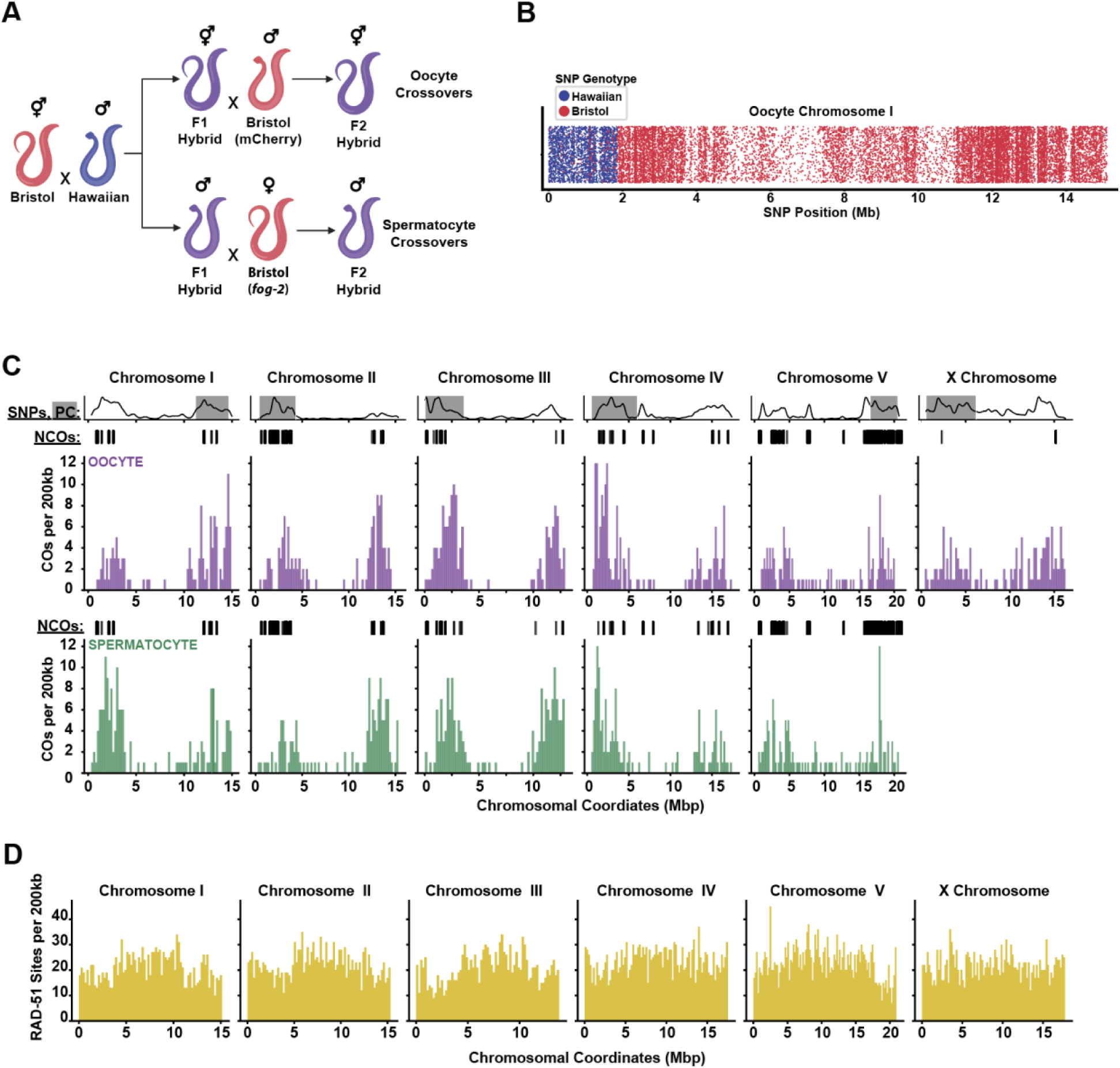
Crossover placement is sexually dimorphic in the C. elegans meiotic recombination landscape. **(A)** The genetic crossing scheme utilized to generate F2 hybrids from mating Bristol and Hawaiian *C. elegans* isolates. F1 hybrids from the first cross were split into two secondary crosses to distinguish oocyte-derived and spermatocyte-derived recombination events. **(B)** A strip plot of a single recombinant chromosome bearing one crossover. Each dot represents individual SNP markers colored by the parental genotype. **(C)** Histograms showing the global distribution of crossovers in oocytes versus oocytes on each chromosome. Crossovers were counted in nonoverlapping 200 kb bins on each chromosome. Above, rug plots showing the spatial distribution of all NCOs detected on each chromosome in each gamete type. Top, a track of line graphs showing the normalized density of SNP markers used in recombination mapping. The location of the pairing centers on each chromosome are shown shaded in grey. Oocyte data is shown in purple and spermatocyte data is shown in green. **(D)** Histograms showing the global distribution of RAD-51 sites, identified from N2 hermaphrodites, in nonoverlapping 200 kb windows.

**Table 1.** Crossovers detected in wild type germ cells.

|  | Chromosome |  |  |  |  |  | Total |
| --- | --- | --- | --- | --- | --- | --- | --- |
|  | I | II | III | IV | V | X |  |
| **Oocyte Crossovers** | 126 | 129 | 144 | 163 | 133 | 142 | 837 |
| **Spermatocyte Crossovers** | 163 | 136 | 162 | 141 | 136 | n.a. | 738 |
| **Mean length of breakpoint interval (bp)** | 1,036 | 1,046 | 1,329 | 1,698 | 2,351 | 2,387 | 1,569 |

### The fine-scale crossover landscapes are sexually dimorphic in *C. elegans* meiosis

To assess whether the global distribution of crossovers was sexually dimorphic on each chromosome, we compared the spatial distribution of crossovers across each chromosome for both spermatocyte and oocyte data. Broadly, both oocyte and spermatocyte crossovers exhibit a bias towards the terminal thirds of all chromosomes. Notably, these results match previous studies describing the distribution pattern for crossovers (53,54). For chromosomes *I*, *II*, and *III*, there appears to be a clearer sex-specific bias for crossover formation on either ‘arm’ of the chromosomes (Figure 1C), with many oocyte crossovers on the same ‘arm’ as the pairing centers (72–74). In contrast to chromosomes *I, II, III,* and *IV*, we detected a greater proportion of crossovers in the central regions of chromosome *V* and the *X* chromosome with a seemingly elevated frequency of central crossovers generally across spermatocyte chromosomes. Using our high-resolution crossover maps, we examined whether any small regions within any of the chromosome arms had elevated crossover formation. Similar to previous studies (54,75), our data did not detect any distinct 1-2 kb regions of extremely elevated crossover rates that resemble recombination hotspots like those seen in mammals and budding yeast (19,44,46). Despite the lack of true recombination hotspots, our data does indicate multiple 200 kb chromosomal regions with a greater density of crossovers, particularly on chromosomes *I, IV,* and *V* in both sexes. (Figure 1C). Thus, while crossover formation is biased towards the chromosome “arms,” we can see fine-scale variability in the crossover landscape suggesting that crossover formation is not random or uniform in these large domains. Notably, the broader trends in our high-resolution crossover data, particularly on chromosomes *I, II,* and *III,* are in agreement with earlier genetic studies assessing sexually dimorphic recombination at megabase-resolution (17,55–57).

We next wanted to more comprehensively analyze the recombination landscape in *C. elegans* by identifying noncrossover (NCO) recombination products in our dataset. To our knowledge, a genome-wide map of noncrossovers in worms has not been published, though prior studies using recombination assays at single loci have demonstrated the frequency and minimal lengths of NCO events at these targeted loci (76,77). In our single-worm WGS data, we developed a method that utilizes the HMM-inferred genotype states at each marker and produced a call set of NCOs based on the theoretical allele frequencies supported by read alignments (See Methods). We were successfully able to identify NCO events on all chromosomes, and the spatial distribution of detectable NCOs appears concentrated in the terminal “arm” domains of each chromosome, with very limited calls in the central regions (Figure 1C, tracks above histograms). With our dataset, we observe that the average number of detectable NCOs per genome is 17-18, and the average minimal length of these detectable NCOs to be approximately 300 bp (Supplemental Figure S3). Notably, the spatial distribution of detectable NCOs largely aligns with regions of exceptional SNP density (Figure 1C, tracks), and the size of the detectable NCOs is approximately the equal to the average spacing of SNP markers genome wide. Thus, although the length and number of NCOs per genome is within prior expectations of the total number of DSBs (76–78), we are likely undercounting the true number of NCO events per genome because of limitations in SNP density and read depth in our dataset. Importantly, there does not appear to be any sexual dimorphisms in the spatial distribution of the detectable NCO events in our dataset (Figure 1C). While it is possible that subtle differences in the distribution of NCO outcomes could eventually be revealed with a greater density of markers and higher sequencing depth, our results here have only detected sexual dimorphisms in the crossover recombination landscape.

To compare our recombination landscapes to the DSB landscape in *C. elegans*, we performed CUT&RUN on the highly conserved recombinase RAD-51, which is a canonical marker of meiotic DSBs in *C. elegans* and marks DSBs in intermediate stages of repair following induction by SPO-11. Cytological analysis of RAD-51 foci along computationally-straightened chromosome axes demonstrates a greater proportion of RAD-51 foci observed on the terminal domains of each chromosome (52). ChIP-seq of RAD-51 in *C. elegans* has also suggested a lack of hot spots (79,80). For CUT&RUN, we detected RAD-51-bound DNA fragments across the genome with base pair resolution using a sliding window approach of read counts in 300 bp bins. Our analysis revealed that while there were no statistically enriched binding sites for RAD-51 (see Methods), the general pattern of RAD-51 bound DNA fragments appears largely random across each chromosome (Figure 1D). Additionally, our method of analysis examining genomic windows in the top 5% for RAD-51 signal indicates that there may be one 300 bp window with higher RAD-51 signal every ten kilobases. In summary, the distribution of DSBs associated with RAD-51 signal appears consistent with the absence of hotspots in the *C. elegans* genome.

### Sexually dimorphic rates of crossing over

Given that the spatial distribution of crossing over appeared sexually dimorphic, we wanted to determine whether the rate of crossing over is also sexually dimorphic across the genome. With our recombination landscape dataset, we first examined the cumulative frequency distributions of crossovers in each sex. The cumulative distributions of crossovers detected on each chromosome illustrate the expected frequency of crossing over between any two loci. We found marked differences between the sexes in crossover frequencies on the arms of chromosomes *I, II*, and *III* (Figure 2A). Spermatocytes display elevated rate of crossover formation in the first five megabases of chromosome *I* (p < 0.001 by KS-test). In comparison, oocytes have higher crossover frequencies between 2.5-5 Mb on the left of chromosomes *II* (p = 0.062) and *III* (p < 0.1, Figure 2A). For chromosomes *IV* and *V*, however, we observed that the frequency of crossover events along their lengths was approximately equal in oocytes and spermatocytes. While there is no recombination on the unpaired X chromosome in *C. elegans* sperm, the rate of crossing over in oocytes is highest on the terminal domains, though this pattern is less pronounced than observations on the autosomes and in concordance with prior genome-wide mapping studies (54). In conclusion, these data show sex biases in the crossover landscape in multi-megabase domains only on chromosomes *I*, *II*, and *III*.

**Figure 2.**
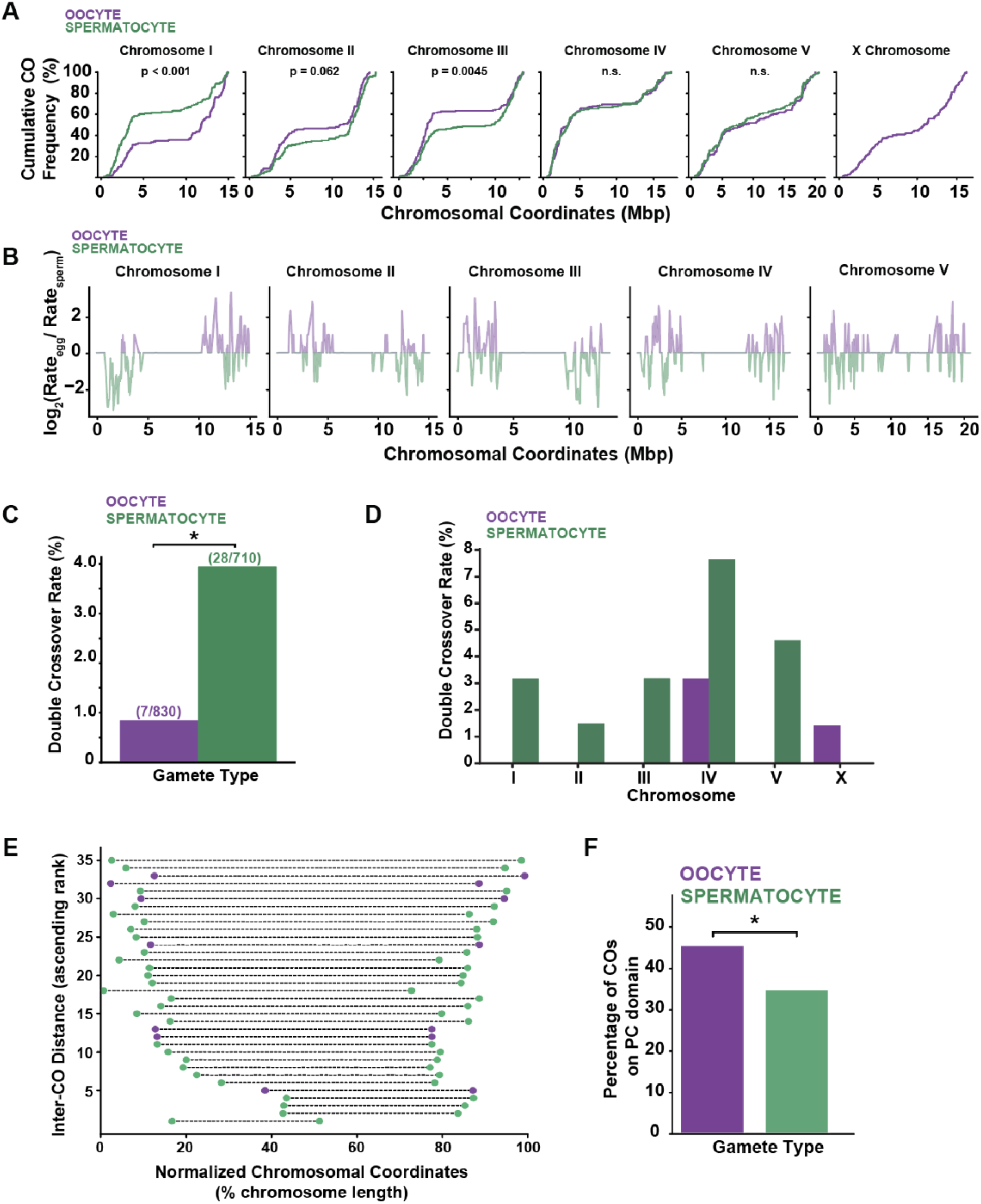
The crossover rate is sexually dimorphic at global, chromosomal, and sub-megabase scales. **(A)** Cumulative frequency plots showing the rate of crossing over across each chromosome length. Significant differences in distributions were determined by KS-test. **(B)** Line plot showing the Log2 values of the ratio of the recombination rate (cM/bp) in eggs versus sperm. Crossover rates were calculated in 200 kb sliding windows with a 50% step size. **(C)** Bar chart showing the global average frequency of double crossovers among all recombinant chromosomes. Asterisk indicates significantly higher DCO rates associated with sperm (p < 0.05 by logistic regression). **(D)** Bar chart showing the frequency of double crossover events on each chromosome in sperm versus eggs. **(E)** Scatter plot showing the pairs of breakpoints for individual chromosomes that experienced a double crossover event. Purple and green dots indicate the double crossover occurred in an oocyte versus spermatocyte genome, respectively. Dashed lines between pairs of dots span the inter-crossover distance normalized by chromosome length. **(F)** A bar chart showing the global frequency of crossovers that occurred on the chromosome “arm” that contains the pairing center. Asterisk indicates a significant difference in association (p < 0.05 by logistic regression) of CO placement on the PC arm between each sex.

Sexual dimorphisms in the crossover frequency at sub-megabase scales could indicate specific regions or features of chromosomes contributing to a sexually dimorphic crossover landscape. To test for sexual dimorphisms in the crossover rate at a finer scale, we performed a sliding window analysis of the crossover rate in 200 kb windows on each chromosome for each sex. Our approach detected many 200 kb regions where each sex has a higher local crossover rate in a non-favored chromosome “arm” (Figure 2B). Consistent with the spatial and frequency distributions of crossovers (Figures 1C and 2A, respectively), we found that chromosomes *I, II,* and *III* have large clusters of 200 kb regions with sexually dimorphic crossover rates (Figure 2B). Despite these large-scale patterns, we found 99 total instances on chromosomes *I* (n=31)*, II* (n=40), and *III* (n=28) where neighboring 200 kb windows display opposite sex biases in the local crossover rate (Figure 2B). Further, while the chromosomal crossover distributions for chromosomes *IV* and *V* are not significantly different between the sexes at the megabase scale, we do see many instances (41 and 64 for chromosomes *IV* and *V*, respectively) where adjacent 200 kb windows have higher crossover rates in the opposite sex (Figure 2B). Taken together, sexual dimorphisms in the crossover landscape persist at sub-megabase scales on each chromosome.

Given that the spatial distribution of crossovers is sexually dimorphic, we then assessed if the overall rate of crossing over also showed sex-specific effects. To further assess sexual dimorphisms in the crossover rate, we calculated the incidence of double crossovers (DCOs) among all recombinant chromosomes at the global and chromosomal scales. The occurrence of DCOs in *C. elegans* is rare and often undetected (53,57–59,81–83). Our data set of 830 oocyte recombinant chromosomes and 710 spermatocyte recombinant chromosomes revealed the global frequency of DCOs is 4.7-fold higher in spermatocytes than oocytes (3.94% spermatocytes vs 0.84% in oocytes; p < 0.05 by logistic regression) (Figure 2C). Notably, we found that this global frequency was not equally shared across all chromosomes or limited to those with sexually dimorphic crossover distributions. In oocytes, we only detected DCOs on chromosomes *IV* (5/168, 3.1%) and the *X* chromosome (2/140, 1.42%), whereas in spermatocytes DCOs were detected on all autosomes. From the analyses of our dataset, we observe that the spermatocyte DCO rate ranges from as low as 1.49% (2/134) on chromosome *II* to as high as 7.63% (10/131) on chromosome *IV*. For spermatocytes, the rate of DCOs on chromosome *IV* is nearly double the global average DCOs frequency in spermatocytes (Figure 2D). For most DCO events, 60% or more of the chromosome length separated the individual CO breakpoints, though some smaller inter-breakpoint distances were observed in spermatocytes (Figure 2E). In summary, the rate of crossing over is elevated in spermatocytes relative to oocytes.

To determine how well the local crossover rates in each sex correlate across many scales, we repeated our sliding window analysis of crossover rates in a range of window sizes from 10 kb up to 2 Mb in 10 kb increments and calculated Kendall’s *tau* as a correlation coefficient (Supplemental Figure S4). For autosomes *I, II, III*, and *IV*, crossover rates show little to no correlation (tau < 0.3) up to window sizes of 30-50 kb. Crossover rates on chromosome *V*, however, remain very weakly correlated up to a window size of approximately 200 kb. At larger window sizes, chromosome *I* is unique in that even in windows of 1 Mb or greater, crossover rates remain only moderately correlated (tau 0.4-0.6). Chromosomes *II, III, IV*, and *V*, in contrast, achieve much higher levels of correlation (tau 0.75-0.9) in megabase-scale windows. Thus, there appears to be fine-scale heterogeneity in the rate of crossing over in spermatocytes versus oocytes at scales of tens to hundreds of kilobases, with some differences persisting up to megabase scale domains. In summary, we find that sexual dimorphisms in the crossover rate are not uniformly shared in their distribution or magnitude across all chromosomes.

We then assessed the relationship between crossing over and proximity to chromosomal pairing centers. Prior studies suggest that crossovers are sometimes bias towards pairing centers (PCs), which are sequence-defined regions on each *C. elegans* chromosome to facilitate homologous chromosome pairing during meiotic prophase I (53–56,72–74,81–83). To assess crossover placement relative to PC-bearing chromosome domains, we defined “arm” and center boundaries independently for each chromosome using a piecewise linear fit to the cumulative crossover frequency distribution, allowing domain boundaries to be determined empirically by the crossover data rather than by fixed physical coordinates (see Methods). Compared to the wild-type spermatocyte crossover landscape, a greater fraction of oocyte crossovers are placed on the PC-bearing “arm” of each chromosome (Figures 1C, 3A). We first wanted to assess whether crossovers are formed within and/or adjacent to clusters of the DNA sequence motifs associated with the pairing center proteins. While we did not observe direct overlaps of crossovers with the sequence motifs themselves, our data revealed that the majority of crossovers are formed approximately 60-70 kb away from clusters of PC sequence motifs (Supplemental Figure S5). Second, we analyzed how frequently crossovers were detected within the multi-megabase PC-bearing “arm” domains on each chromosome (Figure 2F). This analysis found that the genome-wide frequency of crossover placement in proximity PC domains was different between oocytes and spermatocytes (45.8% oocytes vs 34.9% spermatocytes; p < 0.05, logistic regression model). In conclusion, the crossover rate is sexually dimorphic at multiple scales in oocytes versus spermatocytes, and some of this sexual dimorphism can be attributed to crossover formation relative to their distance from the chromosomal pairing centers.

### Normal dosage of synaptonemal complex proteins promote sexually dimorphic crossover placement and constrains crossover rate

Since our analysis of the recombination landscape reveals widespread sexual dimorphism in the placement and rate of crossing over, we wanted to explore the association of crossing over with large-scale chromosome structures in meiosis. Faithful execution of the recombination program depends on the pairing and close juxtaposition of homologous chromosomes that is maintained through synapsis via the synaptonemal complex (SC; (2)). Additionally, decreased abundance of central region SC proteins alters the crossover rate (26,84) and shifts the crossover distribution in a sex-specific manner (38). Specifically, SNP recombination mapping at select loci on Chr. *II* found that heterozygosity for null alleles of the SC central region components *syp-2* and *syp-3* alter sexually dimorphic crossover distribution for Chr. *II* (38). To examine these effects genome-wide, we performed our high-resolution recombination mapping protocol using *C. elegans* that are heterozygous for a null allele of the central region SC component *syp-2.* Briefly, we were able to identify 1,334 crossovers across 1,292 recombinant chromosomes in total for *syp-2 (ok307)/+* progeny (Table 2) in our dataset. Our analysis of the cumulative frequency distributions of crossovers in each wild-type and *syp-2 (ok307)/+* mutant oocytes revealed a statistically significant shift in the crossover landscape on chromosomes *III* (p = 0.0058 by KS-test) and *V* (p = 0.046). Unlike oocytes, we only observed a statistically significant shift in the mutant spermatocyte crossover distribution on chromosome *II* (p = 0.042) (Figure 3A). We then performed pairwise tests comparing the crossover distribution between all four experimental groups. Notably for chromosomes *I* and *II, syp-2 (ok307)/+* mutant sperm display crossover distributions with a greater magnitude of difference relative to wild-type oocytes, indicating that the sexual dimorphism on these chromosomes is exacerbated in mutant sperm. Most notably, when comparing *syp-2 (ok307)/+* mutant oocyte distributions to wild-type spermatocyte distributions, the *syp-2 (ok307)/+* mutant oocyte crossover landscape converged towards the wild-type spermatocyte landscape most chromosomes. Chromosome *I,* however, was the sole exception where a statistically significant difference was maintained between *syp-2/+* mutant oocytes and wild-type spermatocytes (p < 0.001 by KS-test) (Figure 3A), consistent with its distinctive behavior across our other analyses. These results suggest that in wild-type oocytes, the dosage of SYP-2 central region SC proteins are important for establishing sexually dimorphic crossover placement on most chromosomes. Additionally, we analyzed our *syp-2 (ok307)/+* mutant samples to detect noncrossover events. Although we detected slightly more NCOs per sample, the overall distribution appeared highly similar to wild type, with no evidence of sexual dimorphism or *syp-2 (ok307)/+* mutant effects. Taken together, these data support prior hypotheses that dosage of proteins critical for chromosome synapsis regulates crossover placement and contributes to the sexual dimorphic recombination landscape observed in wild-type germ cells.

**Figure 3.**
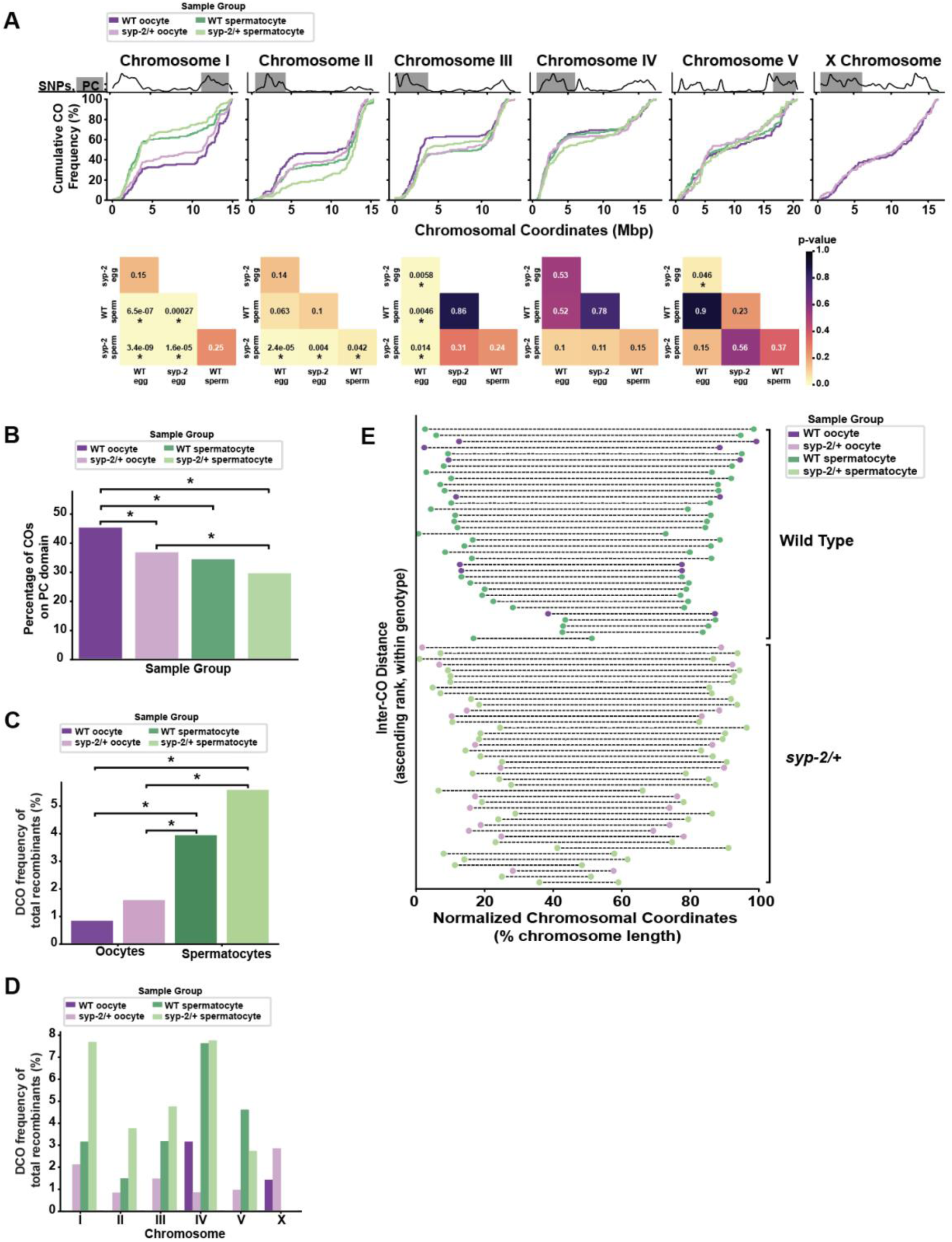
SC protein dosage influences the chromosome-wide crossover distribution and rate. **(A)** Above, cumulative frequency plots showing the rate of crossing over across each chromosome length for wild type and *syp-2/+* heterozygous germ cells. Below, heatmaps of p-values from pairwise KS-tests comparing the cumulative distributions across experimental groups. Asterisks indicate p< 0.05. **(B)** A bar chart showing the global frequency of crossovers that occurred on the chromosome “arm” that contains the pairing center. Asterisk indicates a significant difference in association (p < 0.05 by logistic regression) of CO placement on the PC arm between each sex and genotype. **(C)** Bar chart showing the global frequency of double crossover events among recombinant chromosomes. **(D)** Bar chart showing the individual chromosome frequency of double crossover events among recombinant chromosomes. **(E)** Scatter plot showing the pairs of breakpoints for individual chromosomes that experienced a double crossover event. Purple and green dots indicate the double crossover occurred in an oocyte versus spermatocyte genome, respectively. Dashed lines between pairs of dots span the inter-crossover distance normalized by chromosome length.

**Table 2.** Crossovers detected in *syp-2/+* germ cells.

|  | Chromosome |  |  |  |  |  | Total |
| --- | --- | --- | --- | --- | --- | --- | --- |
|  | I | II | III | IV | V | X |  |
| **Oocyte Crossovers** | 144 | 119 | 137 | 118 | 104 | 144 | 766 |
| **Spermatocyte Crossovers** | 126 | 110 | 132 | 125 | 75 | n.a. | 568 |
| **Mean length of breakpoint interval (bp)** | 1,319 | 1,204 | 1,249 | 1,585 | 1,305 | 2,057 | 1,411 |

We then tested whether the association between crossover placement and the PC-bearing “arm” domain seen in our wild-type dataset was a feature of the *syp-2/+* crossover landscapes. Our wild type data showed that crossovers were spaced approximately 60-70 kb from the nearest cluster of motifs bound by the pairing center proteins. While we did not observe a statistically significant shift from this distance in *syp-2/+* mutant oocytes, *syp-2/+* spermatocytes showed a significant increase in the median distance up to 80 kb (Supplemental Figure S5; p< 0.05 by Kruskal-Wallis H-test and post-hoc Dunn’s test). Further, both types of mutant germ cells displayed a significant genome-wide reduction in the frequency of crossovers placed on the PC-bearing “arm” domain of chromosomes (Figure 3B). Compared to the wild-type oocyte frequency (45.8%), *syp-2/+* oocytes placed 37.7% of their crossovers on the PC-bearing “arm” domains genome wide. In the mutant spermatocytes, while there was a reduction in the frequency of crossovers placed on the PC “arm” domains relative to wild type (30.5% vs 34.9%), this difference was not significantly different. Using a logistic regression model of the frequency of crossovers placed on the PC “arm” domains across experimental groups, the wild-type oocyte frequency was significantly higher than all other groups tested and the mutant oocyte frequency was higher than the mutant spermatocytes (Figure 3B; all p-values < 0.05). In summary, the reduction of SC central region protein SYP-2 appears to weaken the genome-wide association of crossovers placed near the pairing centers in mutant germ cells.

We next wanted to address whether the genome-wide rate of crossing over is also affected in *syp-2 (ok307)/+* mutant germ cells. Similar to what was observed with a cytological crossover marker when *syp-2* was partially knocked down in oocytes (26), the average incidence of DCO events was elevated in both *syp-2 (ok307)/+* mutant spermatocytes and *syp-2 (ok307)/+* mutant oocytes relative to their respective wild-type rates. The genome-wide incidence of DCO chromosomes was observed to be 1.59% in mutant oocytes (∼1.9-fold increase) and 5.58% in mutant spermatocytes (∼1.4-fold increase) (Figure 3C). While there is no significant difference based on genotype differences within each sex, we did observe significant differences in the DCO frequency for all other pairwise comparisons (both p-values < 0.05, logistic regression). When we examined the frequency of DCO events separately for each chromosome, we found elevated DCO formation in both *syp-2 (ok307)/+* mutant oocytes and *syp-2 (ok307)/+* mutant spermatocytes for nearly every chromosome (Figure 3D). These *syp-2/+* mutant oocyte results contrast with wild-type oocytes, where we only observed DCOs on chromosome *IV* and the *X* chromosome. Further, our dataset found that in *syp-2 (ok307)/+* mutant spermatocytes, chromosomes *I* and *II* both displayed greater than two-fold increases in the frequency of DCO events in comparison to wild-type spermatocytes DCO frequencies. Similar to our analysis of the inter-breakpoint distance for DCO chromosomes in wild-type germ cells, our *syp-2/+* mutant data indicate that the majority of DCO breakpoints are still separated by more than half of the length of the chromosome (Figure 3E). However, while all inter-breakpoint distances were greater than 30% of the chromosome length among wild-type DCOs, we did identify 2 spermatocyte DCOs and one oocyte DCO in our *syp-2/+* data with inter-breakpoint distances below 30% of the chromosome length (Figure 3E, bottom). Taken together, our analysis of high-resolution crossover maps in *syp-2/+* mutant germ cells suggest that wild-type sexual dimorphisms in crossover placement and the overall rate of crossing over in both sexes may be at least partially influenced by dosage of the SC central region protein SYP-2.

### Sex-specific crossover associations with germline gene expression

To determine whether any of the sexually dimorphic features of the crossover landscape arise from specific chromosomal features, we first tested the association of the crossover distribution with sequence level genome annotations. We tested for enrichment or depletion in intergenic regions, genes (including sub-gene annotations such as each UTR, exons, introns, and CDS), gene regulatory sequences (transcription factor binding sites, promoters, and enhancers) as well as other sequences such as non-coding RNAs (such as meiotically expressed piRNAs) and transposons (Supplemental Figure S6). In spermatocytes, we did find that crossovers were enriched in “ncRNAs” on chromosomes *I* and *III* (Log_2_(fold) values of 1.58 and 2.77, respectively), which mirrored the sexually dimorphic crossover distributions on these chromosomes. Notably, in the genome annotations we used from Ensembl, “ncRNA” describes a subset of coding genes with non-coding splice variants of their mRNAs. Notably, there was no significant enrichment or depletion for nearly all other features tested. Taken together, these results suggested that a subset of genes with non-coding splice variants in *C. elegans* may drive sex differences in crossover formation on chromosomes *I* and *III*.

To determine if gene expression in the germline shapes crossover distribution, we performed enrichment/depletion analyses only on genes expressed during meiosis using a published RNA-seq dataset that examined transcription levels specifically in the germline of each sex (85). We first labeled our complete set of annotated genes with specificity for oogenesis, spermatogenesis, genes commonly expressed in both germlines, or neither. Labels for shared or sex-specific expression were determined based on the presence of normalized read counts of ≥ 2 in the germline (85). When we tested for fold enrichment or depletion of crossovers overlapping with these subsets of genes, we found that spermatocyte crossovers had significant enrichments in genes expressed in germlines on chromosomes *III* and *V* (Log_2_(fold) values of 0.52 and 0.60, respectively). In contrast, for oocyte crossovers, there were no statistically significant associations with meiotic gene expression states (Figure 4A-B). Overall, these trends suggest that some chromosomal regions actively undergoing transcription may be shaping the crossover landscape differently in each sex.

**Figure 4.**
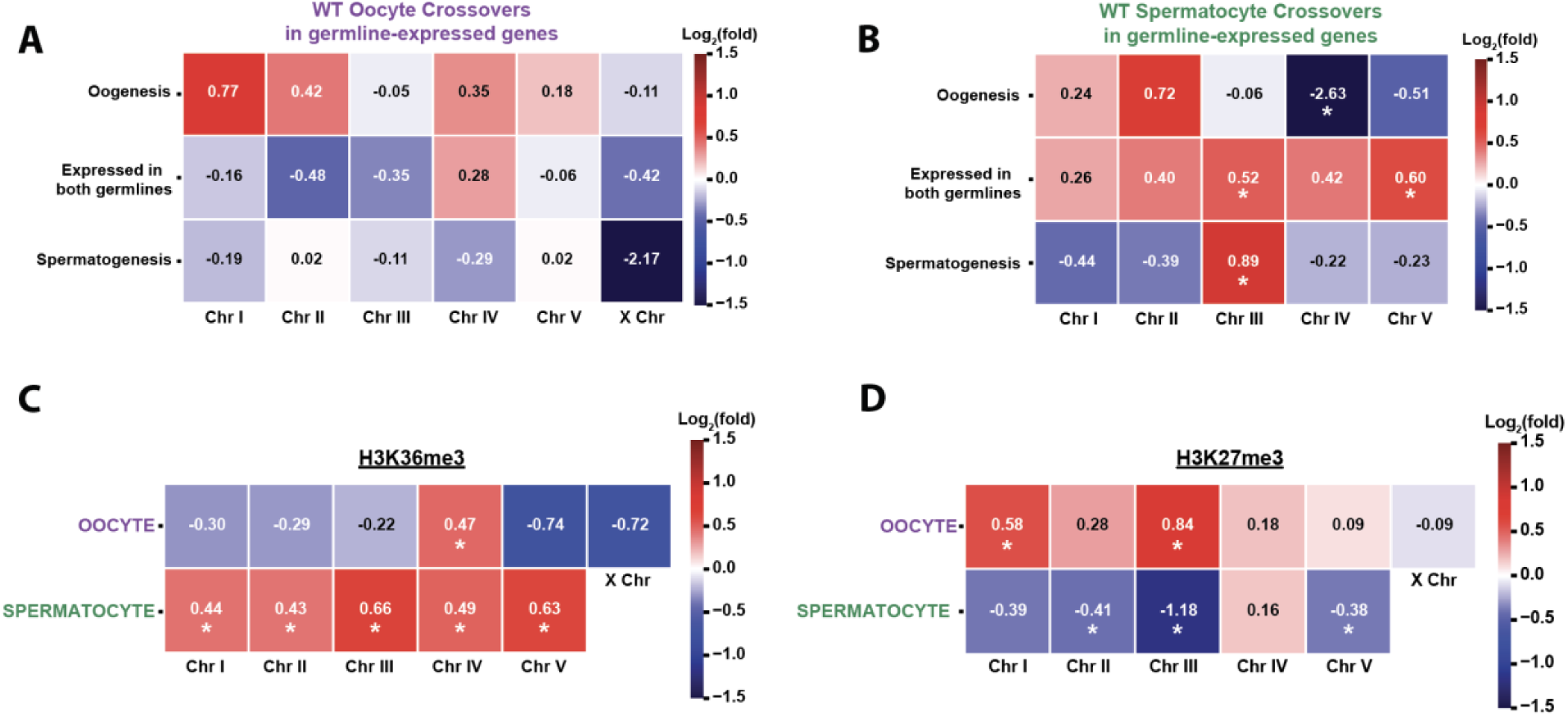
Sexual dimorphisms in crossover placement differentially associate with gene expression and chromatin states in wild-type germ cells. (A) Heatmap showing the log2(fold) association of crossovers in germline expressed genes in wild type oocytes. (B) Heatmap showing the log2(fold) association of crossovers in germline expressed genes in wild type spermatocytes. (C) Heatmap showing the log2(fold) association of wild-type crossovers with H3K36me3 ChIP-seq peaks. (D) Heatmap showing the log2(fold) association of wild-type crossovers with H3K27me3 ChIP-seq peaks. Asterisks indicate p-values < 0.05 by hypergeometric test when compared to simulated null distributions.

### Sex-specific associations of crossovers with chromatin states in wild-type germ cells

Gene expression states and the DSB landscape are influenced by chromatin states in multiple organisms (44,47,48,80,86–89). Since we found an enrichment of crossovers in germline expressed genes within *C. elegans* spermatocytes, we tested the association of crossovers with sex-specific germline landscapes of the euchromatic histone modification H3K36me3 and the heterochromatic histone modification H3K27me3 (66). For the euchromatic modification H3K36me3, spermatocytes displayed significant enrichment of crossovers in euchromatic regions on all five autosomes (Log_2_(fold) values 0.43-0.66, p-values < 0.05; Figure 4C). In contrast, oocytes showed a trend of crossovers depleted in H3K36me3 regions on chromosomes *I, II, III, V,* and *X* (Log_2_(fold) values −0.22 to −0.74). When we compared the difference of fold changes in spermatocytes versus oocytes, chromosomes *I, II, III*, and *V* were all significantly different (p-values < 0.05 by hypergeometric test) (Figure 4C). Only for the euchromatic H3K36me3 landscape on chromosome *IV* did we find that both oocytes and spermatocytes have a significant association with the crossover landscape on chromosome *IV*, (Log_2_(fold) values of 0.47 and 0.49, respectively; p-values < 0.05) (Figure 4C). In summary, we find that crossover formation in spermatocytes, but not oocytes, is strongly associated with chromatin marked by H3K36me3.

We then tested for sexually dimorphic association of crossovers with heterochromatin. For the heterochromatic mark H3K27me3, oocytes displayed an enrichment of crossovers in these regions on chromosomes *I* and *III* (Log_2_(fold) values of 0.58 and 0.84, respectively; Figure 4D). In contrast, spermatocyte crossovers were depleted from heterochromatic regions on chromosomes *II, III*, and *V* (Log_2_(fold) values of −0.38 to −1.18; Figure 4D). For chromosomes *I, II*, and *III*, the chromosomes with the most sexually dimorphic crossover landscapes, we found the difference in fold enrichments between the sexes to be statistically significant (all p-values < 0.05 by hypergeometric test). Overall, our results suggest that the sex-specific crossover landscapes in *C. elegans* may be also influenced by local differences in chromatin structure and accessibility near sites of active transcription in the germline.

## Discussion

### Homologous chromosome pairing and sexual dimorphisms in crossover distribution

Our studies report the sexually dimorphic global recombination map for *C. elegans* meiosis. In our dataset, we find a greater proportion of crossovers on the pairing centers “arms” of chromosomes in oocytes (Figure 2). Notably, chromosomes *I, II*, and *III* display the greatest magnitude of sexual dimorphisms for crossover distribution relative to pairing center “arms.” The chromosome-specific magnitude of these sexual dimorphisms may reflect the intrinsic differences in chromosome architecture and domain organization, consistent with emerging evidence from *Drosophila* that chromosome-specific properties broadly influence recombination landscape outcomes (90). Consistent with this, these same chromosomes also have the greatest number of pairing center motif clusters relative to their total length (74). Based on our data and published data from others, we hypothesize that the pairing of homologous chromosomes could play a role in shaping the wild-type sexual dimorphisms of the crossover landscape in each sex. One proposed model for the progression of recombination describes how all DSB sites are competing for the accumulation of pro-crossover factors to determine the crossover versus non-crossover outcome (91–94). If homolog pairing enables faithful crossover formation, then DSB sites in early-paired regions of chromosomes might have a temporal advantage in processing recombination intermediates, which increases the likelihood to form crossovers compared to DSB sites with less time in alignment. In comparison to spermatogenesis, the total duration of meiotic prophase I, as well as time spent in the window for chromosome pairing, is nearly twice as long in *C. elegans* oogenesis (95). Recent studies demonstrate that the timing differences inherent to *C. elegans* spermatogenesis versus oogenesis are not solely causal for differences in the observable rate/number of crossover events(64), yet it remains possible that this temporal factor may still partially affect the spatial distribution of crossover events on some chromosomes. Thus, the prolonged duration of pairing in wild-type *C. elegans* oogenesis may one factor that contributes to the broad-scale differences in the crossover distribution on the pairing center “arms” of these chromosomes.

### Chromosome synapsis and sexually dimorphic crossover distributions

In many species, synapsed homologous chromosomes are required for recombination and accurate chromosome segregation (96). In *C. elegans,* growing evidence suggests that both the proper abundance and the spatial arrangement of SC proteins regulate crossover number (26,38,97,98). Our study finds that SYP-2 dosage shifts the spatial patterning of crossovers. While we observe sexual dimorphisms in crossover placement on chromosomes *I, II*, and *III* in wild-type germ cells, we find that the crossover landscape in *syp-2/+* mutant oocytes shifts to a landscape that appears highly similar to the landscape seen in wild-type sperm. How might the synaptonemal complex establish the normal, sex-specific crossover distribution in oocytes? Notably, a previous *C. elegans* study found that *syp-2/+* oocytes display SC discontinuities, delayed completion of synapsis into mid-pachytene, and shifts in the crossover distribution on one chromosome that was assessed. Further, these observations and their effect sizes are weaker or not apparent in spermatocytes (38). The stage at which these SC assembly defects occur coincides with the loss of signal for some PC proteins in wild-type hermaphrodite germlines (73). The convergence of *syp-2/+* mutant oocytes to the distribution seen in wild-type spermatocytes (crossovers further away from early-paired chromosome ends) could potentially be explained by these synapsis defects. Mutant *syp-2/+* oocytes in mid-pachytene may have altered or weakened homolog connectivity, which could lead to DSBs across the chromosome becoming more equally competitive for acquiring pro-crossover factors for recombination processing. Notably, the central region proteins of the synaptonemal complex (SYPs) help drive the localization of pro-CO proteins to interhomolog recombination events (28,99). Additionally, there is emerging evidence that SC proteins may be facilitating signal transduction and movement of proteins (*e.g.* ZHP-3) that promote crossover formation (100). We therefore speculate that any favorable effects of the early-paired chromosome domains on recombination processivity in oocytes may be lost when synapsis is fragmented, unstable, and/or delayed along the length of some chromosomes, thereby shifting the overall distributions away from the PC domains. We note that Chromosome *I* was the sole exception to the convergence of mutant oocyte crossover distributions towards those in wild-type spermatocyte, maintaining a significant difference between mutant oocytes and wild-type spermatocyte crossover distributions. Whether this reflects the same genetic incompatibility affecting chromosome *I* crossover counts in our wild-type oocyte data, or a chromosome *I*-specific feature of SC dynamics remains unclear. Future studies could clarify how SYP-2 dosage and SC integrity impart sexual dimorphisms in crossover placement, and how this impacts specific chromosomes, in wild-type germ cells.

### Sexually dimorphic crossover rates

Our dataset observes that all autosomes in spermatogenesis undergo higher double crossover events rates than oogenesis. The genome-wide frequency of DCO chromosomes is 4.7-fold higher in spermatocytes than oocytes, and DCO events were detected on all wild-type spermatocytes autosomes but only chromosomes *IV* and the *X* chromosome in oocytes. These results extend prior genetic interval-based studies that detected elevated DCO frequencies in *C. elegans* spermatogenesis to a genome-wide scale at higher resolution (55,60,61). Sexual dimorphisms in double crossover rates have been documented in other species as well, namely in humans and some plants (101–105). Further, our analysis of *syp-2/+* heterozygous germ cells demonstrates that a lower dosage of critical SC central region proteins not only shifts the spatial distribution of crossovers but elevates the incidence of double crossover chromosomes in both sexes genome-wide. These data are consistent with previous findings indicating that the mechanism(s) of crossover frequencies could be affected by SYP protein dosage or discontinuous synapsis (25,26,28,84,106,107). While we do not have the statistical power to confidently calculate crossover interference in this study, our results support a hypothesis that the overall distribution and rate of crossing over between the sexes could also be a product of differences in the strength of crossover interference. Multiple models have been proposed with regards to effectors of interference (5,25,26,94), however, the precise molecular mechanisms that lead to this phenomenon remains unclear. Further research analyzing a much greater number double crossover events, or experimental approaches that enhance high-throughput collection of DCO data, on all chromosomes in *C. elegans* will more precisely characterize sex-and/or chromosome-specific variations in the strength of interference.

### Crossovers in euchromatin versus heterochromatin

For oogenesis, we find that the crossing over is enriched in H3K27me3 marked heterochromatic regions on chromosomes *I* and *III* (Figure 4). In contrast in spermatogenesis, we note a negative association of crossovers with H3K27me3 and an enrichment of crossovers in H3K36me3 euchromatic regions. These results suggest that not only do sex-specific chromatin distributions influence the crossover landscape in each sex, but that the *C. elegans* recombination program in each sex may be differentially affected by the presence of different chromatin modifications. Given that recombination intermediate processing is reliant on the recruitment and accumulation of multiple pro-crossover factors at DSB sites (8,99,108–114), the processing of recombination intermediates may be kinetically unfavored in heterochromatin where the more tightly compacted DNA may be inherently more difficult for the recruitment and accumulation of pro-crossover factors. Alternatively, the prolonged duration of Prophase I in oocytes could be amenable to the slower kinetics of DSB repair in heterochromatin. Despite this potential kinetic delay, there are known mechanisms for DSB processing in heterochromatin that support a model for the initiation and maturation of oocyte crossovers in these dense chromatin states. Studies in Drosophila demonstrate that DSB sites are relocated outside of heterochromatic compartments and processed for repair near the nuclear envelope in a more physically accessible environment (115,116). Further, DSBs made in euchromatin versus heterochromatin are processed with similar kinetics, indicating that heterochromatic DSBs are not wholly refractory to crossover formation (117).

Our analysis of the overlap of oocyte crossovers in heterochromatin could be due to the maintenance and/or presence of these chromatin modifications during DSB processing and crossover formation. In *C. elegans*, it is still unknown whether chromatin marks are dynamically remodeled around sites of active recombination or whether recombination intermediates relocate to more repair-permissive chromatin environments. In either case, these potential mechanisms could promote sex differences in recombination. For spermatocytes, crossover events could preferentially take place in regions of accessible chromatin that happen to coincide with transcription. Future studies could elucidate why spermatocytes and oocytes differ in their crossover placement relative to different chromatin modifications.

### Sexually dimorphic crossover distributions and gene evolution

Our results indicate that crossover recombination in *C. elegans* spermatogenesis displays a greater association with genes that are actively expressed during in meiosis. Recombination in genes can be mutagenic (118) and/or introduce genetic diversity in progeny through reciprocal exchange genetic content on homologs. Further, studies have found that transposable element activity, large scale differences in DNA methylation, and spermatocyte-specific euchromatic chromatin modifications create a “promiscuous” state that likely enhances the evolution of current and novel genes (119,120). Thus, our detection of elevated crossover formation within gene bodies during spermatogenesis support hypotheses that suggest meiotic genes are rapidly evolving (121,122). Notably, the ‘out of testis’ hypothesis poses that male meioses may be a unique source of selection on reproductive genes (120). Our data aligns with these hypotheses and evidence from Drosophila studies that demonstrated genes expressed in spermatogenesis are under positive selection, which promotes adaptation and fixation of beneficial alleles in populations (123,124). The exact molecular mechanism driving this spermatocyte-specific preference for crossovers in euchromatin remains elusive, and further investigation is warranted to understand the rate at which recombination in spermatocytes may be promoting the evolution of meiotic genes in *C. elegans* populations.

## Materials and Methods

### *Caenorhabditis elegans* strains and maintenance

All strains were incubated at 20°C and maintained on nematode growth medium (NGM) plates seeded with the OP50 strain of *Escherichia coli.* Strains used in this experiment include the following: N2 (wildtype from Bristol, England), CB4856 (wildtype from Hawaii, United States), EG7841 (oxTi302 [eft-3p::mCherry::tbb-2 3’UTR + Cbr-unc-119(+)] I), and CB4108 (*fog-2*(q71) V), and DLW188 (*syp-2*(ok307)/tmC16 [*unc-60*(tmIs1210)] V). All genetic crosses were done by mating L4 stage males and hermaphrodites on NGM plates and screening for cross progeny after 3-4 days.

### Crossing schemes for sex-specific crossover mapping

For wild-type recombination mapping, Parent (P0) N2 hermaphrodites were mated to CB4856 males to generate Bristol/Hawaiian F1 hybrids. Individual F1 progeny were placed onto their own plates and separated into two separate cross schemes. To assess oocyte recombination, F1 hermaphrodites at the L4 stage were then mated to EG7841 males, and F2 cross progeny marked with red fluorescent bodies were collected and briefly transferred to a new plate. To assess spermatocyte recombination, F1 males at the L4 stage were mated to CB4108 hermaphrodites, and male F2 cross progeny were collected and briefly transferred to a new plate before sucrose floatation. For recombination mapping in the *syp-2/+* heterozygote background, non-Unc GFP-positive hermaphrodites of the DLW188 strain were mated to CB4856 males to generate F1 hybrids. F1 hybrids that are GFP-negative (lacking the balancer) are heterozygous null mutants for the *syp-2* gene, and these progenies were selected for mating and production of the F2 generation that is sequenced. As in the wild-type crosses, *syp-2/+* F1 hybrids were mated to either EG7841 males or CB4108 hermaphrodites to distinguish oocyte versus spermatocyte-derived recombination.

### Sucrose floatation and isolation of individual F2 progeny

To minimize bacterial contamination in downstream gDNA sample preps, we performed sucrose floatation on pooled worms from each cross scheme, respectively. Previously collected worms were immediately washed from plates with 8mL cold M9 buffer and transferred to 15mL glass centrifuge tubes using a glass Pasteur pipette. Collected worms were centrifuged at 3000rpm at 4°C and washed in 4mL of fresh M9 twice. To separate worms from bacteria and other debris, 4mL of 60% sucrose solution was very gently added to 4mL of M9 buffer and worms, preserving the distinct layers of sucrose and M9. The mixture was then spun at 5000 rpm at 4°C for 5 minutes. Using a glass pipette, the floating layer of worms were transferred to a new glass centrifuge tube on ice and brought up to 4mL in fresh M9. Worms were then incubated at room temp for 30 minutes and gently vortexed every 5 minutes. Worms were washed three times in equal volume of fresh M9 were performed before storing collected worms at −80°C in 1.5mL microcentrifuge tubes containing M9 buffer. To isolate individual F2 progeny before sequencing, 10-20 worms were transferred via glass Pasteur pipette into M9 buffer on a glass well slide. Individual worms were then pipetted into separate microcentrifuge tubes with approximately 10 microliters of M9. gDNA isolations were performed using the Invitrogen PureLink™ Genomic DNA Mini Kits and eluted with a total volume of 40 microliters. Individual worms’ gDNA samples were then transferred into a single well of a 96-well PCR plate before library prep. Plates containing F2 progeny were then briefly stored at −80°C.

### Illumina Whole Genome Sequencing, marker selection, and data processing

Our method for the high throughput analysis of meiotic recombination in individual genomes of *C. elegans* is achievable with as little as 0.75-1.0 total nanograms of genomic DNA collected per individual. We sequenced 610 individual wild-type genomes (300 oocyte-derived samples and 310 spermatocyte-derived samples) and 600 genomes (300 oocyte-derived samples and 300 spermatocyte-derived samples) from the *syp-2* heterozygous null crosses at nearly half of the cost compared to available commercial methods. For Illumina short-read sequencing, library preparation was performed on individual worms for by the University of Oregon’s Genomics and Cell Characterization Core Facility. The short-read libraries were then sequenced on an Illumina Novaseq (2 x 150 bp). The average read depth was approximately 10X for our wild-type samples and 5X for our mutant samples (Supplemental Figure S1).

Previously, our lab sequenced and assembled genomes *de novo* for the N2 Bristol and CB4856 Hawaiian strains (67). The SNP sites chosen for recombination mapping in this study were filtered from the parental VCF to only include homozygous sites (mean variant allele fraction >99.3%) and sites that did not overlap with repeats, low-complexity sequences, and transposons identified by RepeatMasker. This leaves a marker set of 213,591 higher quality SNP markers across the genome. We improved upon previous SNP maps to cover more than 99.9% of each chromosome’s length with 99.97% of the total genome covered. The average SNP spacing on each autosome (*I, II, III, IV*, and V) is 645 bp, 354 bp, 580 bp, 674 bp, and 262 bp, and the average SNP spacing on the *X* chromosome is 1020 bp. Many regions, such as the terminal “arm” domains of each chromosome have a much greater density of SNPs, and thus resolution of crossover detection is high. The centers of each chromosome have a much lower density of markers, with the largest gap between SNPs being 106kb on chromosome *V*.

Illumina short reads from each individual F2 genome were trimmed using Trimmomatic (125) to remove adapter and barcode sequences. The trimmed reads were then aligned to our lab’s previously generated N2 Bristol reference genome (NCBI accession number PRJNA907379; (67). All resulting variant positions comparing our N2 Bristol and CB4856 Hawaiian genomes are in relation to this N2 Bristol assembly. Aligned reads in SAM format were then sorted using SAMtools (126) and converted to BAM files. Using Picard, read groups were added via AddOrReplaceReadGroups, and duplicate reads were filtered using MarkDuplicates. BAM files with filtered duplicate reads (127) were used to call variant and homozygous reference sites using GATK HaplotypeCaller (128) to generate per-sample VCF files. Each sample’s VCF file was then filtered to only include the higher quality SNP sites suitable for recombination mapping.

### Reconstruction of F1 chromosomal products of meiosis

To aid crossover detection, a Hidden Markov Model approach was used to reconstruct the chromosomes present after crossing over between the Bristol and Hawaiian homologous chromosomes in oocytes and spermatocytes of the F1 generation. A previously developed pipeline, TIGER (68), was adapted for use on our samples because of its high accuracy at calling crossovers at sparse sequencing coverage (97.5% accuracy at 0.1X and 99.3% at 10X). Control for running each TIGER script was combined into a single shell script. Read counts and alleles for each SNP marker were generated from the output of GATK’s HaplotypeCaller, and input files were prepared as described in (Rowan et al. 2015; https://github.com/betharowan/TIGER_Scripts-for-distribution). To briefly review their framework, the ratio of read counts between the N2 Bristol and CB4856 Hawaiian backgrounds are transformed into a discrete alphabet and 6 states represented as AA, AU, AB, BU, BB, and UU. AA represents a homozygous N2 Bristol state, AB is the heterozygous state, U represents uncertainty in the homozygous state, and UU represents no reads/information at a given marker. A marker site with a minimum of 5 total reads where 100% of the reads are from a single parental background are required to assign genotypes AA or BB. If there are fewer than five reads aligned or both parental alleles were observed at the same marker, genotypes were inferred from a multinomial distribution. Observing reads in a homozygous parental background should follow a binomial distribution such that 99% of reads (accounting for 1% sequencing errors) are from either N2 or CB4856, and 50% for the heterozygous state. Genotypes were then assigned according to the maximum value between the homozygous parental and heterozygous probabilities. Genotypes AU or BU were assigned unless there was an equal probability of each genotypic background at these markers with low coverage and/or uncertain parental origin.

An HMM is then generated for each individual genome to infer Bristol or Hawaiian haplotype blocks on each chromosome. Transmission and emission probabilities are estimated per sample for each model, and hidden states (homozygous N2, homozygous CB4856, or heterozygous) were then determined via the Viterbi algorithm. Output files for each sample that contain the intervals of each inferred hidden state were then converted to Pandas data frames in Python for further processing.

### Feature engineering for Random Forest Classification

Random Forest Classification is a supervised machine learning method that can be used to automate the classification of chromosomes as recombinant or parental using a model trained on numerical features of each sample’s WGS data. Our random forest classifier distinguishes transitions between each haplotype block on a single chromosome as Crossover or “not crossover”. Each haplotype interval is given as a pair of SNP coordinates denoting the start and end of each N2 or CB4856 interval. Between each haplotype interval, a new “transition” interval was created that is defined as two SNP markers whereby the start of the transition interval is the last SNP in a haplotype interval prior to transition, and the end of the transition interval is the first SNP in the following haplotype interval. To label these transition intervals as potential crossovers, numerical features associated with these transitions must be made, and these numerical data were calculated in varying window sizes from 10kb up to 10Mb centered on each transition interval. First, for each window size, the percentage of SNPs with CB4856 Hawaiian alleles upstream and downstream were calculated, and then the difference between these values was recorded as a separate feature. For an ideal crossover breakpoint, a transition interval will have the highest possible difference in the percentage of CB4856 alleles upstream vs downstream. Secondly, for each window size, the cumulative distribution of both CB4856 and N2 alleles, respectively, was stored as an array such that each marker’s index in the array contains the cumulative percentage of that given genotype up to that marker’s position in the window. A third array was then generated as the difference between the CB4856 cumulative distribution array and the N2 array. Then, the values for the difference in cumulative genotype distributions for the marker positions at the start of the transition interval and the marker at its end were averaged to assign an overall difference in cumulative genotype distributions to the transition interval. For an ideal crossover breakpoint, a transition interval will have the highest possible difference between each genotype’s cumulative distribution in that window. All these calculations were performed in windows rather than across the whole chromosome to increase the sensitivity of detection for both single and double crossover events on a single chromosome. Finally, in each individual chromosome, all numerical features associated with each transition interval were converted to a zero to one scale by normalizing to the maximum value of each feature across all transitions.

### Training and Random Forest Classification of Crossover Sites

The training data for this classifier includes 217 of the 3,660 chromosomes sequenced from our wild-type crossing scheme. The training data includes a nearly even mixture of spermatocyte and oocyte samples as well as a mixture of non-recombinant, single crossover and double crossover chromosomes determined by manual visualization and confirmation. The classifier was set up as a forest of 500 decision trees, and the model was trained on 70% of the training dataset and then validated on the remaining 30%. Validation showed that the model had 99.7% accuracy in appropriately labeling crossover versus non-crossover transition intervals. The most features that provided the greatest amount of accuracy to the model were calculations performed in window sizes of 500kb, 1MB, and 2MB. Transition intervals labeled as crossover breakpoints were retained for further analyses Chromosomes were visually inspected to further validate automatic classification of crossover breakpoints. Notably, the first 50kb of the X chromosome was responsible for 40 samples having improper crossover calls in our wild-type dataset. In each of these samples’ X chromosomes, there were two crossovers identified: one in the 50kb region and another crossover as close as 1Mb away. Because three SNP intervals were improperly identified as crossovers in all X chromosomes in these 40 samples, these markers/intervals were then removed from classification in the final crossover datasets (wild-type and *syp-2/+*) given the extremely unlikely probability of such a high rate of crossing over in the same narrow region. In our wild-type dataset, 18 crossovers on 17 chromosomes were recovered by visualization and manual inspection of the highest probable transition intervals. In our mutant dataset, 86 crossovers on 79 chromosomes were recovered by manual inspection of the highest probably transition intervals. Finally, 13 wild-type samples and 36 mutant samples did not have sufficient read coverage (much lower than 1X) and/or high noise leading to unresolvable crossover breakpoints.

### Identification of noncrossover recombination products

Our strategy for the identification of noncrossover products relies on the interpretation of allele frequencies within short haplotype blocks determined by each sample’s HMM via the TIGER protocol (68). First, we restricted our marker set further to only include sites where the mapping quality was a minimum of 40 to reduce the likelihood of false-positives due to mapping artifacts. We anticipated two types of noncrossover products based on our crossing schemes: (1) regions where read alignments indicate a Hawaiian allele frequency of approximately 50% that is surrounded by N2 Bristol genomic background and (2) regions where read alignments indicate homozygous N2 Bristol sequence surrounded by Bristol/Hawaiian heterozygosity. Given the small size of reported noncrossover products in *C. elegans*, Arabidopsis, and mice (76,129,130), we restricted our search to putative noncrossover intervals that were less than one kilobase in size. Additionally, within each interval, we required that there be a minimum of two SNP markers and at least two reads across each marker site. For type 1 noncrossover products, we required the Hawaiian allele/read frequency to be between 40-60% of the total reads. For type 2 noncrossover products, we required a minimum of 90% Bristol allele frequency to allow for one mis-mapped read in our higher coverage samples while precluding noncrossover calls in our samples with less than 10X read coverage. For each putative noncrossovers interval, the longest consecutive series of SNP markers whose HMM-inferred states match our type-1 versus type-2 expectations and meet the criteria listed above were retained in our final call set.

### Assessing the RAD-51-bound DSB landscape via CUT&RUN

The RAD-51 genomic landscape was defined using CUT & RUN (131) on N2 hermaphrodites. Moderate changes were made to the existing protocol to optimize this for the DNA repair protein, RAD-51. Adult hermaphrodites, were collected using a sex-specific size exclusion protocol. Worms were flash frozen and stored at −80 ℃ until further use. A 100 ul pellet of worms was used for this protocol. To disrupt the cuticle and permeabilize the nuclei, worms were bead-beaten with ZR Bashing Tubes (Zymo Research) for 20 minutes and subsequently transferred to a new tube. CUTANA™ Concanavalin A Conjugated Paramagnetic Beads (EpiCypher) were added to bind to the membrane of the nuclei. Following the addition of the beads, a RAD-51 rabbit antibody made by the Libuda lab was used to target RAD-51-bound DNA double-strand breaks. The remaining sample preparation and DNA extraction was completed as stated by Emerson & Lee, 2022. DNA fragments isolated by CUT & RUN underwent a DNA fragment analysis to confirm enrichment. DNA fragments were submitted to the University of Oregon Genomics and Cell Characterization Core for Illumina library preparation and paired end Illumina Sequencing on the NovaSeq6000. Three no-antibody control and three antibody-treated samples were sequenced with a minimum sequencing depth of 10 million reads per sample.

To determine the binding sites of RAD-51 across the genome, we adapted the DESeq2 workflow (132) and carried out analyses using PyDESeq2 (133). Although this software is largely used for differential expression analysis of RNA-seq data, it’s read count modeling approaches are suitable for other genomic analysis like ChIP-seq, CUT&RUN, and other chromatin-profiling methods. Rather than modeling counts across genomic intervals associated with coding regions, we used a sliding window approach to generate 300 bp windows (with 150 bp overlaps/step size) and counted aligned reads in each window. Following the adjustment of p-values to correct for false discovery rate, we found no 300 bp windows to be enriched for RAD-51 binding. This is, in part, due to the high-burden of the number of initial tests computed (>666,000 windows) and the relatively low signal-to-noise ratio across windows. Visual inspection showed a nearly uniform distribution of RAD-51 sites across the genome, in line with expectations from a lack of recombination hotspots in *C. elegans* (54). We then sorted our data by Log2 fold-change values and retained windows in the top 5% as a heuristic proxy for 300 bp regions with much higher RAD-51 signal.

### Genomic features correlated with crossing over

Gene annotations (gene, mRNA, exon, CDS, etc.) were downloaded from Ensembl for the ce11 genome assembly. Annotations for regulatory sequences, transcription factor binding sites, etc, were downloaded from Wormbase for the ce11 assembly as well. The coordinates for these sequence level annotations were then remapped onto our N2 Bristol genome assembly using Minimap2 alignments in the LiftOff program (134,135). Sex-specific germline RNA-seq annotations (85) were taken and applied to our sequence level “gene” annotations in the remapped Ensembl dataset. Sex-specific germline ChIP-seq data (NCBI BioProject PRJNA475794) were aligned to the N2 genome and peaks were called using MACS3 (136) using parameters as previously described (Tabuchi et al. 2018).

To identify pairing center regions to test direct overlap, PC motifs from Phillips et al. 2009 were identified and cluster analysis was performed in our genome assembly using MCAST (137) with maximum motif spacing of 100 bp to be considered in the same cluster. The distance from each crossover to the nearest motif cluster was calculated via the ‘closest’ function in BedTools (138). Statistically significant differences in this distance metric were calculated by using the Kruskal-Wallis H-test followed by post-hoc Dunn’s test to determine the significance of pairwise comparisons across experimental groups. In measuring the frequency of crossovers in the larger pairing center domains, we analyzed overlaps across the chromosome ‘arm’ where 70% of PC motifs reside.

To define chromosome “arm” domains that contain the pairing center for each chromosome, we applied a piecewise linear regression to the cumulative crossover frequency distribution along each chromosome using the *pwlf* python package (https://github.com/cjekel/piecewise_linear_fit_py). Two breakpoints were identified as the positions at which the slope of the cumulative crossover distribution changed significantly, demarcating transitions between low-recombination central regions and high-recombination terminal arm domains. Domain boundaries were fit independently for each chromosome and each sex, and the resulting arm definitions were used to assess the frequency of crossover placement within PC-bearing versus non-PC-bearing arms.

To determine the fold enrichment (or depletion) of crossovers in meiotically-expressed genes and germline-specific chromatin states, the Genomic Association Tester (GAT, Heger et al. 2013) was used to simulate a null distribution of overlaps from 10,000 randomly shuffled intervals per annotation type. Significance of fold change (observed vs null/expected) was determined by hypergeometric test. Significance of the difference between fold changes in each sex was determined by examining the fold change ratio of oocytes vs spermatocytes via hypergeometric test.

## Acknowledgements

We thank N. Kurhanewicz, C. Cahoon, E. Toraason, A. DiNardo, J. Brown, and C. Crahan___ as prior/current members of the Libuda Lab for constructed strains, thoughtful meetings, discussion, and/or comments during manuscript preparation. We are extremely grateful to the University of Oregon Genomics and Cell Characterization Core Facility for the abundance of library prep and sequencing performed across >1200 samples. Additionally, we are grateful to the University of Oregon’s Research Advanced Computing Services for enabling our analyses on Talapas, our campus HPC cluster (URL: https://racs.uoregon.edu). Finally, we thank the Caenorhabditis Genetics Center for providing multiple strains for this study (funded by National Institutes of Health P40 OD010440).

## Funding

This work was supported by National Institutes of Health T32HD007348 to Z.D.B. and H.R.W., and National Institutes of Health R35GM128890 and University of Oregon start-up funds to D.E.L.

## Data availability

Genomic sequence data for our lab’s parental N2 and CB4856 strains were previously submitted to the NCBI BioProject database (NCBI accession number PRJNA907379). Illumina sequencing data produced for this study can be found under the accession number [in progress; will be provided prior to/at the time of publication].

## Competing Interests

The authors declare that no competing interests exist.

---

**Supplemental Figure S1.**
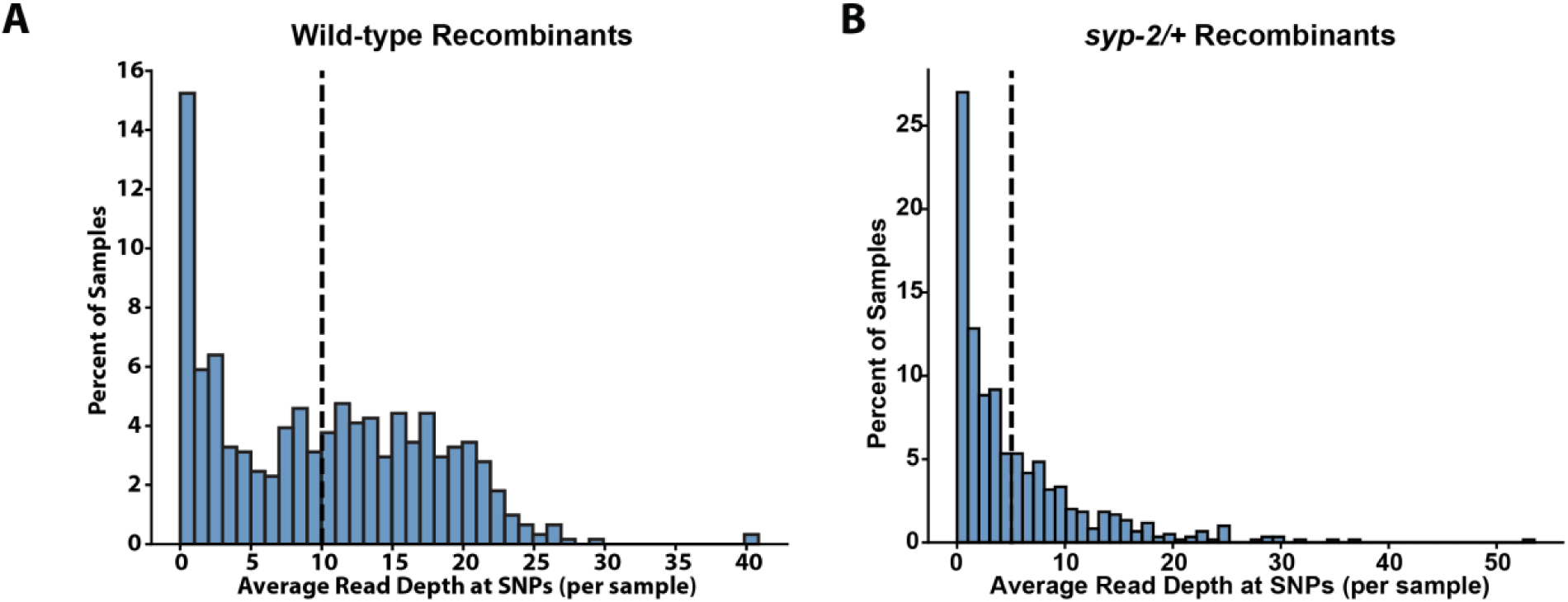
Sequencing read depth of F2 recombinants. **(A)** A histogram depicting the distribution of read coverage at marker sites across all individual wild type samples. **(B)** A histogram depicting the distribution of read coverage at marker sites across all individual *syp-2/+* samples. Vertical dashed line indicates the average across all samples.

**Supplemental Figure S2.**
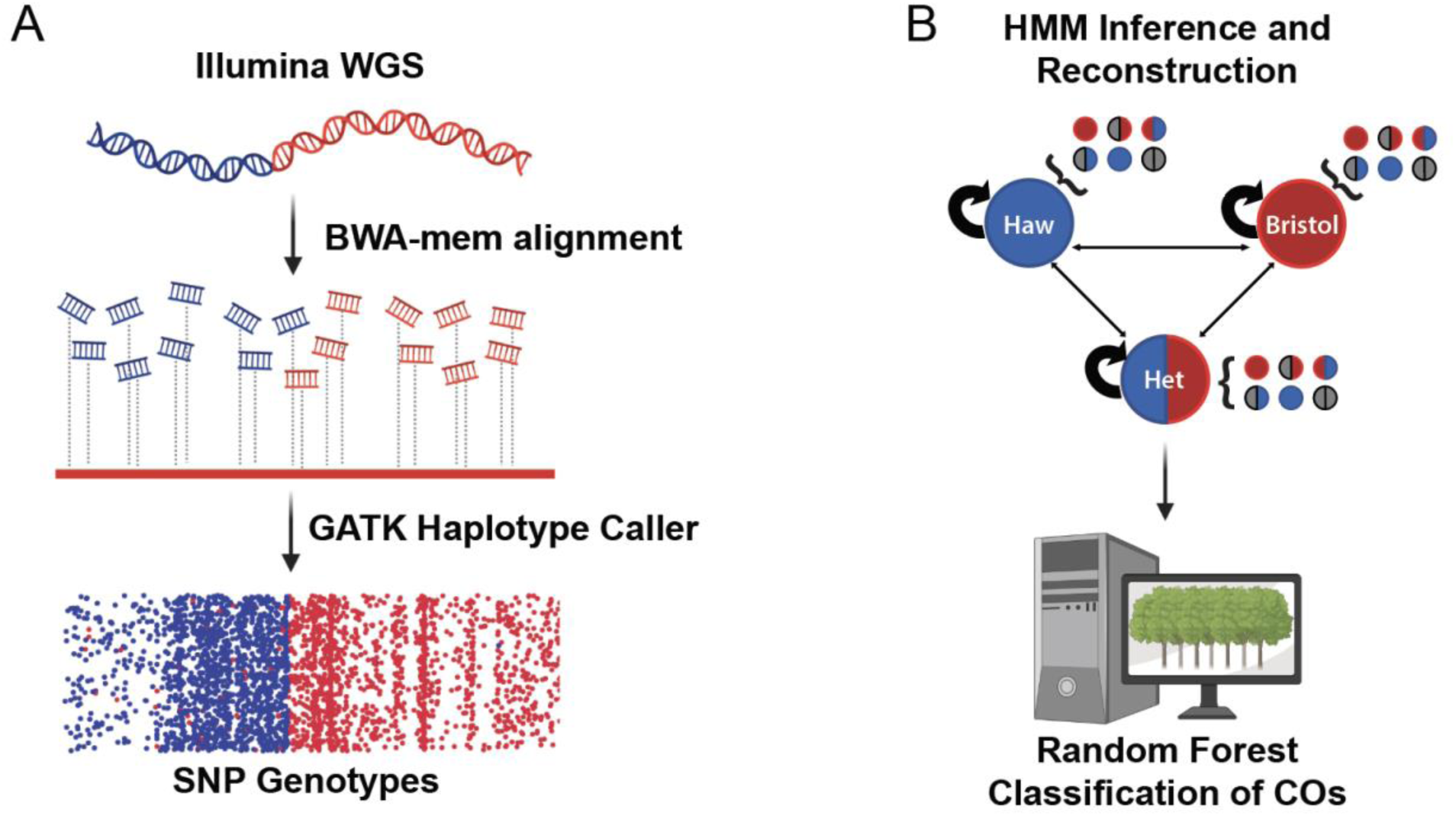
Computational pipeline for processing of sequencing data. **(A)** Data collection and pre-processing including Illumina whole-genome sequencing, read alignment, and SNP calling. **(B)** Schematic of Hidden Markov Model based reconstruction of recombinant chromosomes and subsequent crossover detection via supervised machine learning and random forest classification.

**Supplemental Figure S3.**
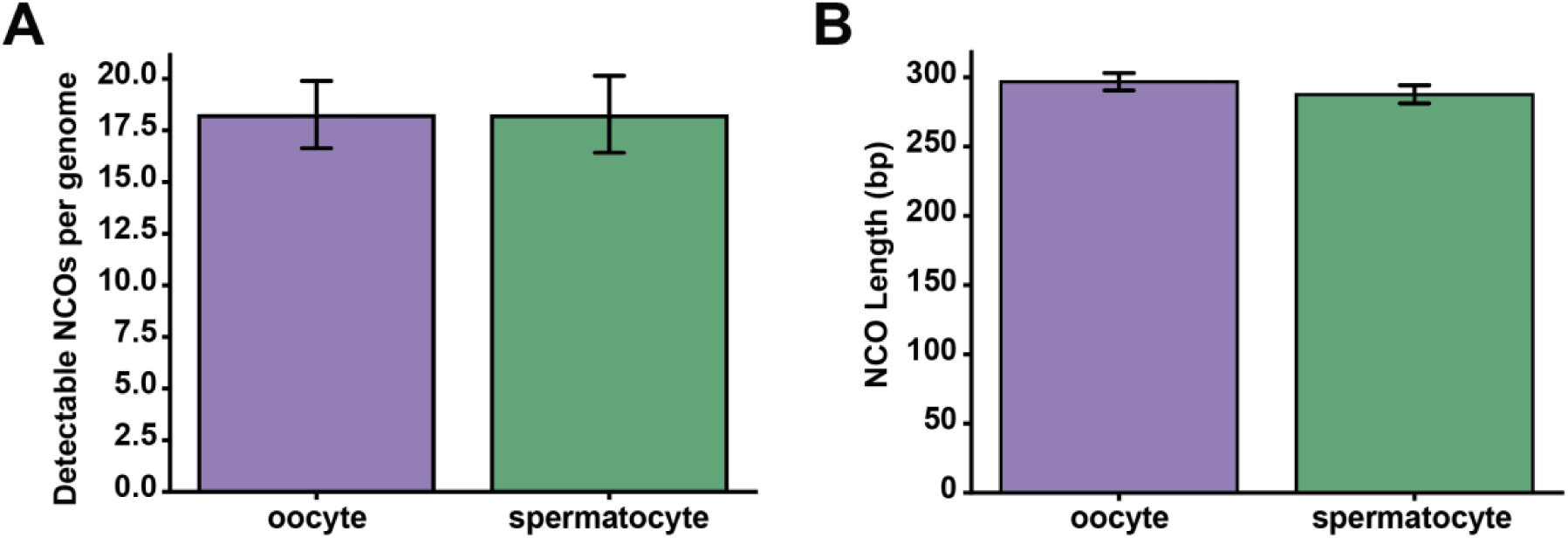
Characteristics of detectable NCO events in wild-type germ cells. **(A)** A bar chart indicating the number of noncrossovers (NCOs) identified in individual oocyte and spermatocyte genomes. **(B)** A bar chart showing the average length of NCO tracts in individual oocyte and spermatocyte genomes.

**Supplemental Figure S4.**
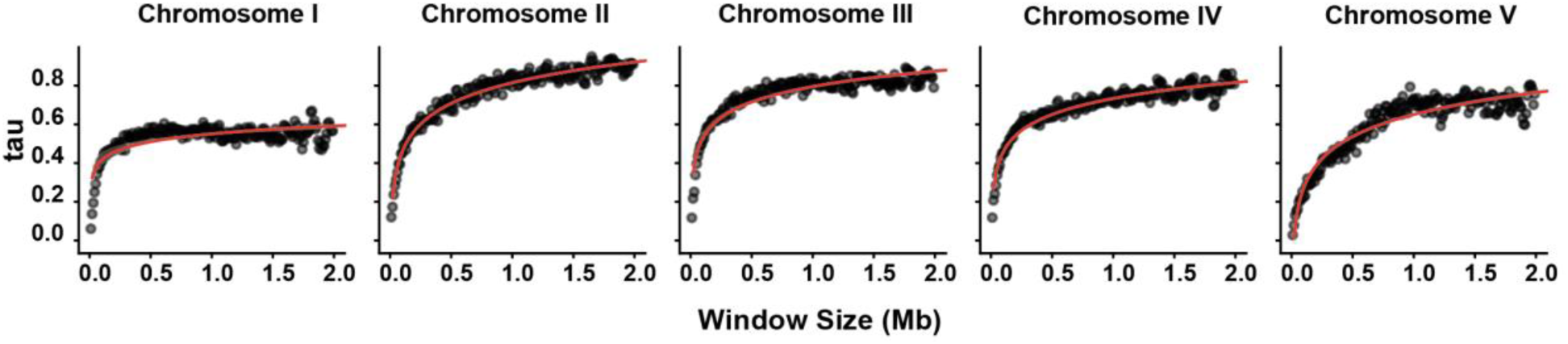
Correlation of egg and sperm crossover rates at multiple scales. Scatterplot showing the correlation of oocyte and spermatocyte recombination rates in sliding windows of varying sizes. Kendall’s tau was calculated as the correlation coefficient, all p-values < 0.05 except 10 kb window size on chromosome *V*. Red line indicates the curve of best fit for each chromosome.

**Supplemental Figure S5.**
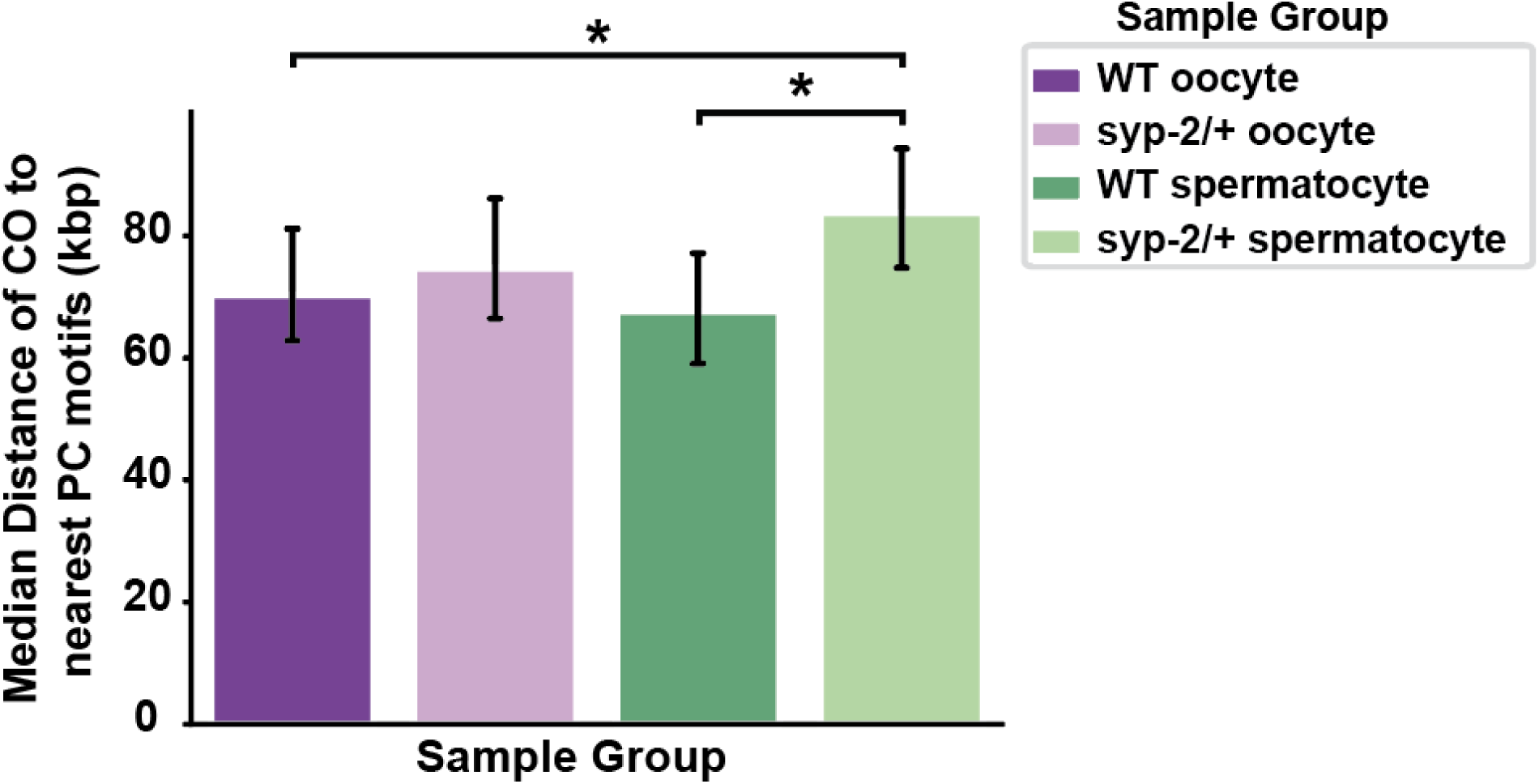
Wild type and *syp-2/+* crossovers associated with chromosomal pairing domains. A bar chart showing the median distance from every detected crossover to the nearest cluster of DNA motifs bound by pairing center proteins. Asterisks indicate p< 0.05 by Kruskal-Wallis H-test followed by post-hoc Dunn’s test.

**Supplemental Figure 6.**
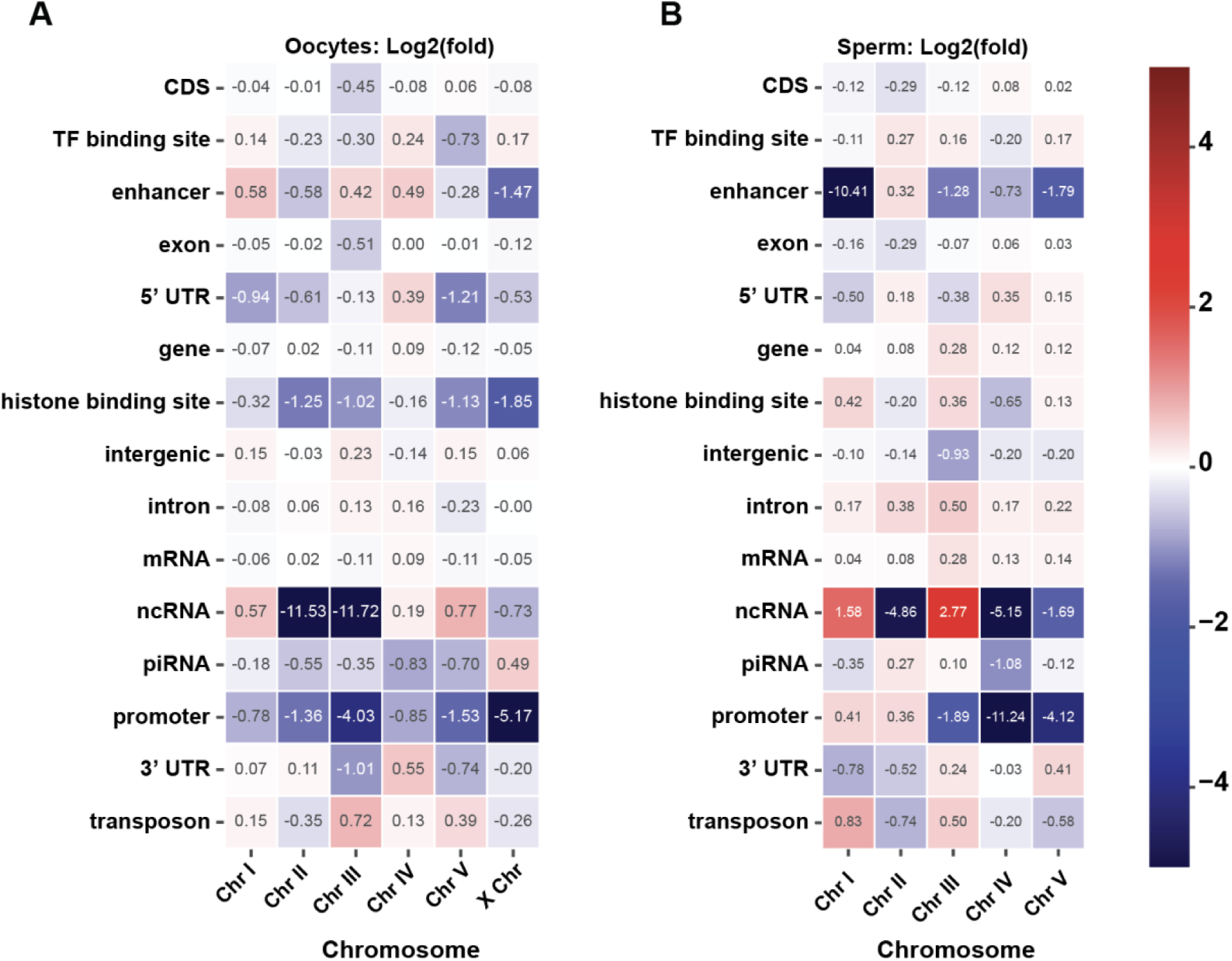
Association of crossovers with sequence-level annotations. **(A)** Heatmap showing the log2(fold) enrichment or depletion of oocyte crossovers with each sequence annotation. **(B)** Heatmap showing the log2(fold) enrichment or depletion of spermatocyte crossovers with each sequence annotation. All annotations were taken from the Ensembl ce11 genome annotation set and remapped to Libuda N2 Bristol genome assembly via LiftOff (See Methods).

## Notes

### Competing Interest Statement

The authors have declared no competing interest.

